# Differences in the path to exit the ribosome across the three domains of life

**DOI:** 10.1101/357970

**Authors:** Khanh Dao Duc, Sanjit S. Batra, Nicholas Bhattacharya, Jamie H. D. Cate, Yun S. Song

## Abstract

Recent advances in biological imaging have led to a surge of fine-resolution structures of the ribosome from diverse organisms. Comparing these structures, especially the exit tunnel, to characterize the key similarities and differences across species is essential for various important applications, such as designing antibiotic drugs and understanding the intricate details of translation dynamics. Here, we compile and compare 20 fine-resolution cryo-EM and X-ray crystallography structures of the ribosome recently obtained from all three domains of life (bacteria, archaea and eukarya). We first show that a hierarchical clustering of tunnel shapes closely reflects the species phylogeny. Then, by analyzing the ribosomal RNAs and proteins localized near the tunnel, we explain the observed geometric variations and show direct association between the conservations of the geometry, structure, and sequence. We find that the tunnel is more conserved in its upper part, from the polypeptide transferase center to the constriction site. In the lower part, tunnels are significantly narrower in eukaryotes than in bacteria, and we provide evidence for the existence of a second constriction site in eukaryotic tunnels. We also show that ribosomal RNA and protein sequences are more likely to be conserved closer to the tunnel, as is the presence of positively charged amino acids. Overall, our comparative analysis shows how the geometric and biophysical properties of the exit tunnel play an important role in ensuring proper transit of the nascent polypeptide chain, and may explain the differences observed in several co-translational processes across species.

## INTRODUCTION

Ribosomes are the key actors of mRNA translation, a fundamental biological process underlying all forms of life. While decoding the mRNA nucleotides into their associated polypeptide sequence, ribosomes regulate the dynamics of translation and other central co-translational processes such as the translocation to cell membranes or protein folding [1, 2, 3]. These processes rely on the structural properties of the ribosome, through interactions with different elements such as binding factors, tRNAs, or the nascent polypeptide chain. For example, it has been shown that some specific sequence motifs associated with antibiotic treatment could stall the ribosome and subsequently arrest translation [4, 5, 6, 7]. This phenomenon is caused by interactions between the ribosome and the nascent polypeptide chain itself: prior to leaving the ribosome, nascent polypeptides first pass through a structure called the ribosome exit tunnel, spanning from the peptidyl-transferase center (PTC) to the surface of the ribosome. As the tunnel can accommodate about 40 amino acids or so [8], its geometry and biophysical properties potentially impact the translation dynamics [9].

Being a functionally important structure, the ribosome exit tunnel needs to be well conserved across species to ensure proper translation of mRNA sequences. On the other hand, the selectivity of arrest sequences to specific species [6, 8], or differences of translational and co-translational mechanisms between eukarya and bacteria [10, 11, 12, 13], for example, suggest that important variations of the exit tunnel structure exist. As such variations have potentially important consequences on the regulation of translation or antibiotic resistance, it is thus crucial to identify and catalog these differences, and more generally understand the evolution of the ribosome exit tunnel.

For the past several years, the ribosome exit tunnel has been the object of intense study to unravel characteristic features, such as its geometric or electrodynamic properties, solvent behavior, rigidity, and dynamic domains [14, 15, 16, 17, 18, 19], to name just a few. As an essential molecular machine present in all living systems, the ribosome has also been extensively used in the past to elucidate phylogenetic relationships via sequence analysis [20]. More recently, several studies have shed light on the relation between the evolution of the ribosome and its function [21, 22]. Specifically, by taking advantage of the sequence information and availability of 3D structures from X-ray crystallography and cryo-electron microscopy (cryo-EM), it has been shown how the evolution of ribosomal RNA (rRNA) has been locally constrained at the beginning of the tunnel around the PTC, where amino acids coded by the mRNA sequence get added to the polypeptide chain. Over the past few years an increasing number of new ribosome structures has been obtained at an atomic resolution of a few Angströms (see below). Hence, it is now possible to extend our understanding of the relation between the biophysical structure of the entire exit tunnel and its evolution across many different species, thereby unraveling local specificities of the tunnel function.

In the present work, we provide a quantitative analysis and comparison of the ribosome exit tunnel structure across a diverse set of species. We compile and compare 20 recently obtained fine-resolution ribosome structures, coming from all three domains of life (bacteria, archae, and eukarya). Upon extracting the coordinates of the tunnels from these structures, we investigate the relation between the geometry of the tunnel and the evolution of the ribosomal structure and its constituent sequences. To achieve this, we introduce and apply a suite of computational methods to study the geometric properties of the tunnel, the local structure of the ribosome near the tunnel, and the conservation of rRNA and ribosomal protein sequences. Our comparative analysis highlights the essential role of the geometric and biophysical properties of the exit tunnel in ensuring proper transit of the nascent polypeptide chain, while revealing potential specificities of the tunnel function across different species.

## RESULTS

### Extraction of the ribosome exit tunnel structure

We compiled and analyzed publicly available ribosome structures obtained from cryo-EM and X-ray crystallography (see Methods and Table 1). These recent structures (9 published in 2017) accounted for 20 different organisms, including 2 archaea, 2 organelles, 8 bacteria and 8 eukaryotes. Most of these structures (16 out of 20) came from cryo-EM maps, with an average resolution of 3.74 Å; the X-Ray structures had an average resolution of 2.92 Å. Replicates of *E. coli*, *T. thermophilus* and *H. sapiens* ribosome structures were also included in our analysis to examine the robustness of our results (see Supporting Information). For each structure, we applied a procedure described in Methods and illustrated in Figure 1, to extract the ribosome exit tunnel geometry, encoded as a set of coordinates describing the trajectory of the centerline and the tunnel radius at each point of the centerline. Upon obtaining these data, we checked the quality of each tunnel reconstruction by comparing the density-fitting score (see Methods) between residues located close to the tunnel and the rest of the structure. All the structures in our dataset showed an increase of the density-fitting score for tunnel residues, suggesting the good quality of maps in the tunnel region.

**Figure 1.**
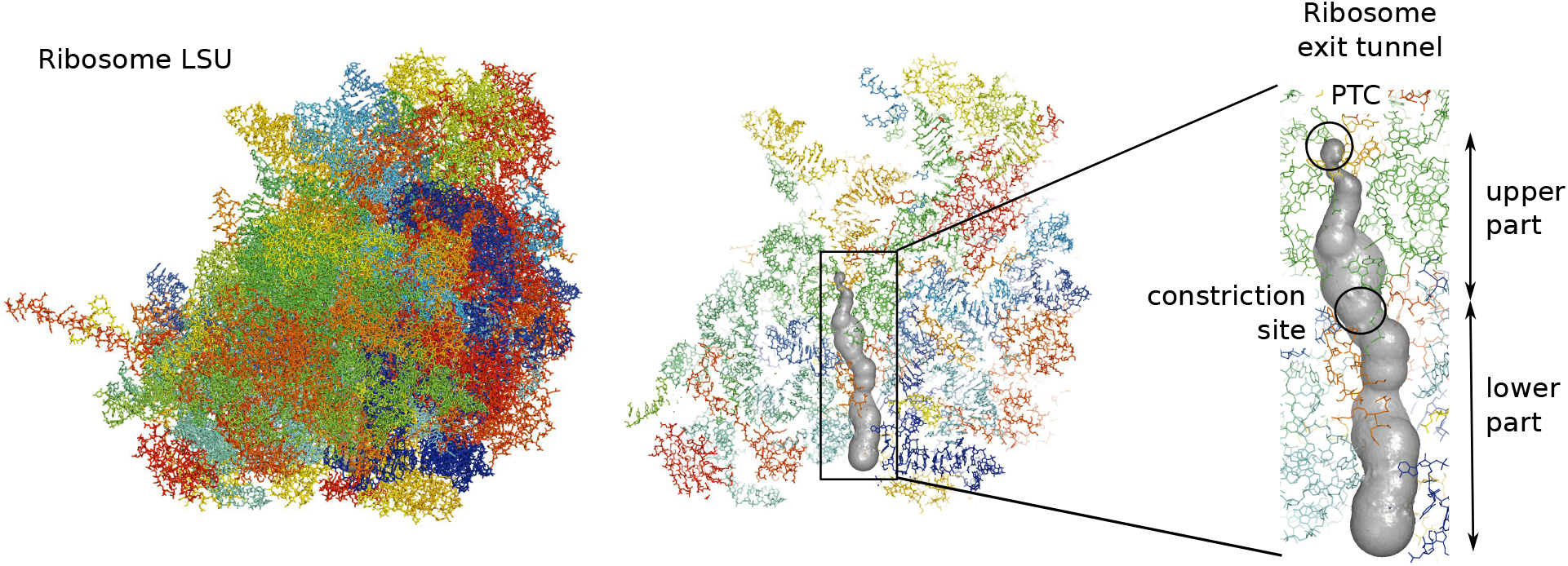
Extraction of the ribosome exit tunnel coordinates. For a given structure of the ribosome large subunit (LSU) represented in the left panel (from Schmidt *et al.* [96]), we first locate the polypeptide transferase center (PTC) and then apply a tunnel search algorithm [76] to reconstruct the geometry of the exit tunnel, illustrated on the right (more details in the Methods section). The inset in the right panel shows the exit tunnel with the PTC and the constriction site that separates the tunnel between its upper and lower parts.

**Table 1.** Ribosomes structures used in our study. First column contains the species, with the domain they belong to (b: bacteria, a: archaea, e: eukarya). The upper part of the table contains the main structures of the study. Replicates for *H. sapiens*, *E. coli* and *T. thermophilus* are in the lower part.

| Species/Organelles | Resolution | Reference | PDB codename |
| --- | --- | --- | --- |
| *Chloroplast (Spinacia)* | 3.8 Å (EM) | Ahmed *et al.* (2017) [81] | 5X8T |
| *Mitochondria (H. Sapiens)* | 3.1 Å (EM) | Amunts *et al.* (2015) [82] | 3J9M |
| *B. subtilis* (b) | 3.8 Å (EM) | Beckert *et al.* (2017) [83] | 5NJT |
| *E. coli* (b) | 2.9 Å (EM) | Fischer *et al.* (2015) [84] | 5AFI |
| *D. radiodurans* (b) | 3.4 Å (X-Ray) | Krupkin *et al.* (2016) [85] | 5JVG |
| *L. lactis* (b) | 5.6 Å (EM) | Franken *et al.* (2017) [86] | 5MYJ |
| *M. smegmatis* (b) | 3.2 Å (EM) | Hentschel *et al.* (2017) [87] | 5O60 |
| *M. tuberculosis* (b) | 3.4 Å (EM) | Yang *et al.* (2017) [88] | 5V7Q |
| *S. aureus* (b) | 3.4 Å (X-Ray) | Matzov *et al.* (2017) [89] | 5NRG |
| *T. thermophilus* (b) | 2.5 Å (X-Ray) | Polinakov *et al.* (2015) [90] | 4Y4P |
| *H. marismortui* (a) | 2.4 Å (X-Ray) | Gabdulkhakov *et al.* (2013) [91] | 4V9F |
| *P. furiosus* (a) | 6.6 Å (EM) | Armache *et al.* (2012) [92] | 4V6U |
| *H. sapiens* (e) | 2.9 Å (EM) | Natchiar *et al.* (2017) [93] | 6EK0 |
| *L. donovani* (e) | 2.9 Å (EM) | Zhang *et al.* (2016) [94] | 5T2A |
| *P. falciparum* (e) | 3.2 Å (EM) | Wong *et al.* (2014) [95] | 3J79 |
| *S. cerevisiae* (e) | 3.9 Å (EM) | Schmidt *et al.* (2016) [96] | 5GAK |
| *T. aestivum* (e) | 5.5 Å (EM) | Gogala *et al.* (2014) [97] | 4V7E |
| *T. cruzi* (e) | 2.5 Å (EM) | Liu *et al.* (2016) [98] | 5T5H |
| *T. gondii* (e) | 3.2 Å (EM) | Li *et al.* (2017) [24] | 5XXB |
| *T. vaginalis* (e) | 3.4 Å (EM) | Li *et al.* (2017) [24] | 5XY3 |
| *E. coli* (b) | 3.9 Å (EM) | Arenz *et al.* (2014) [99] | 3J7Z |
| *T. thermophilus* (b) | 2.8 Å (X-Ray) | Osterman *et al.* (2017) [100] | 5VP2 |
| *H. sapiens* (e) | 3.6 Å (EM) | Khatter *et al.* (2015) [101] | 4UG0 |

### Analysis of global features indicates larger bacterial tunnels and more variation in the lower part

To compare the geometry of the ribosome exit tunnel across different species, we first examined global geometric features, such as the length, tunnel-wide average radius, and volume (see Supplementary Data for specific values). The range of values for the tunnel length (88.8 *±* 6.0 Å) and that for average radius (5.4 *±* 0.4 Å) were both consistent with previous observations [8, 23]. Upon ordering the species by their exit tunnel volume (see Figure 2), we found a perfect separation between bacteria and eukaryotes, with archaea in between. Specifically, bacterial tunnels (including the ones from organelles) are larger than eukaryotic ones, with mean (3.85 *±* 0.37) *×* 10^4^ Å^3^ for bacteria, compared to (2.78 *±* 0.13) *×* 10^4^ Å^3^ for eukaryotes. We similarly analyzed the length and average radius (Figure 2) to see if the volume variation could be mainly explained by one of these two variables. We obtained a less clear separation among the three domains of life, but still observed a similar trend for the length (91.6 *±* 3.6 Å for bacteria and 83.3 *±* 2.9 Å for eukarya) and average radius (5.7 *±* 0.3 Å for bacteria and 5.1 *±* 0.1 Å for eukarya), suggesting that both contribute to the observed difference in volume.

**Figure 2.**
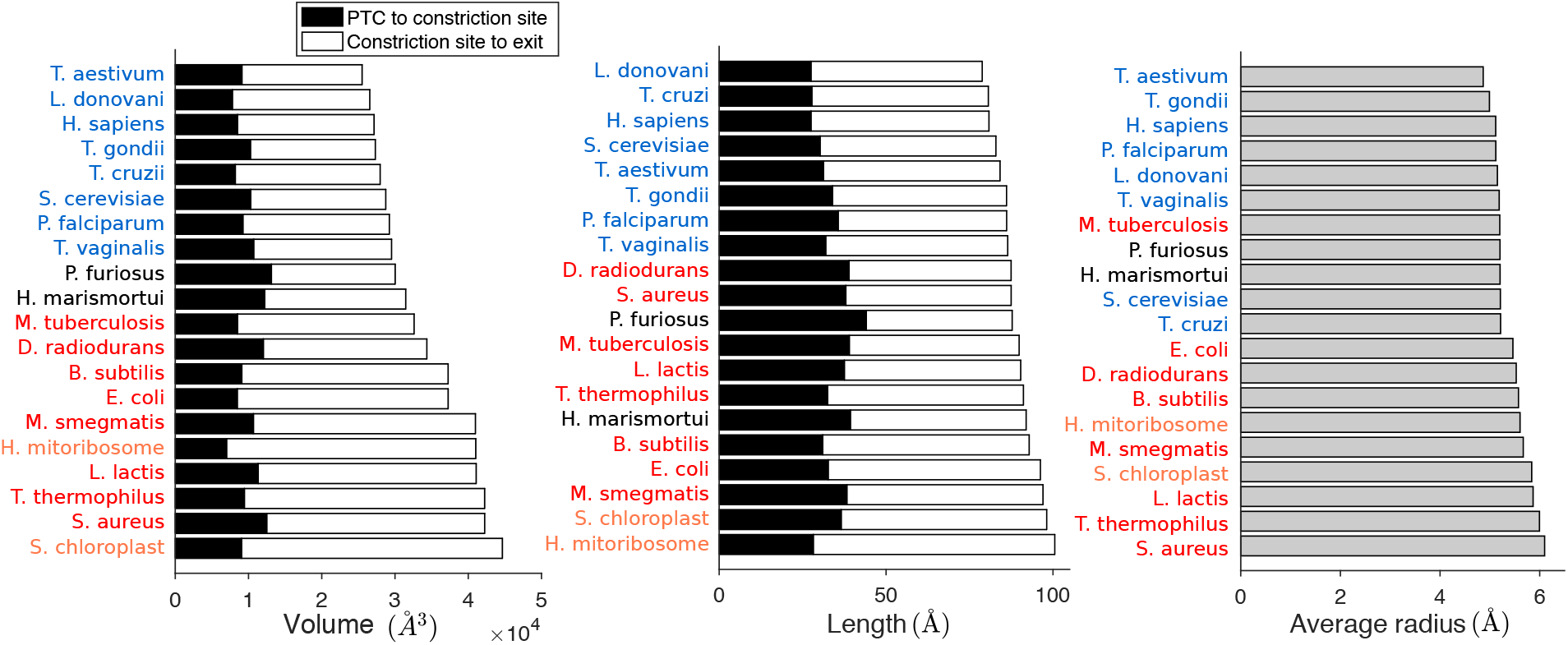
Volume, length and average radius of the ribosome exit tunnel across different species. Horizontal bar plots represent the ordered volume (left), length (middle) and average radius (right) of the tunnels from our dataset. The volume and length are decomposed into two subparts, separated by the constriction site (see Methods). Species are specified and colored by their respective domains (plus organelles, in orange): bacteria (red), archaea (black), eukarya (blue).

To study how these geometric features vary along the tunnel, we carried out a more refined analysis by partitioning the tunnel into two sub-parts separated by the “constriction site” (see Figure 1), a conserved central region constricted by the uL4 and uL22 protein loops. Upon locating the constriction site (see Methods) for each tunnel, which were at position 34.1 *±* 4.8 Å from the start of the tunnel (see Figure 2), we studied how the volumes in the upper (from start to constriction site) and lower (from constriction site to exit) parts respectively correlated with the volume of the entire tunnel. We found a strong correlation for the lower part (Pearson correlation *r*^2^ = 0.9649, p-value *p <* 10^−4^), while no significant correlation was observed for the upper part (*r* ^2^= 0.085), suggesting that the upper part of the tunnel is geometrically quite conserved and most of the variation across species comes from the lower part. The total length also shows a better correlation with the length of the lower part of the tunnel (*r*^2^ = 0.72, *p <* 10^−3^, compared with *r*^2^ = 0.32, *p <* 10^−3^ for the upper part). While the upper part accounts for 38.4 *±* 5.0% of the total tunnel length, it captures only 30.0 *±* 6.9% of the total tunnel volume, suggesting an increase of the average radius in the lower part. Computing the average radius in the two regions indeed showed an increase from 4.7 *±* 0.3 Å in the upper part to 5.8 *±* 0.6 Å in the lower part.

### Hierarchical clustering of tunnel radius variation plots reflects the species phylogeny

As we found domain-specific geometric variations of the tunnel that are more amplified in the lower part of the tunnel, we aimed to quantify these variations more precisely. We first examined the 3D coordinates of the tunnel centerline and found that 96.7 *±* 1.0% of their variations could be explained by fitting to a straight line (see Supplementary Data). We therefore simply parametrized the centerline of the tunnel by its arc length and studied the associated radii (see Methods), leading to a radius variation plot for each species (Figure 3A and Figure S1). To compare these plots, we introduced a distance function (see Methods) and evaluated it for each pair of species. The resulting pairwise distance matrix was then used to cluster the species, yielding the hierarchical tree shown in Figure 3B.

**Figure 3.**
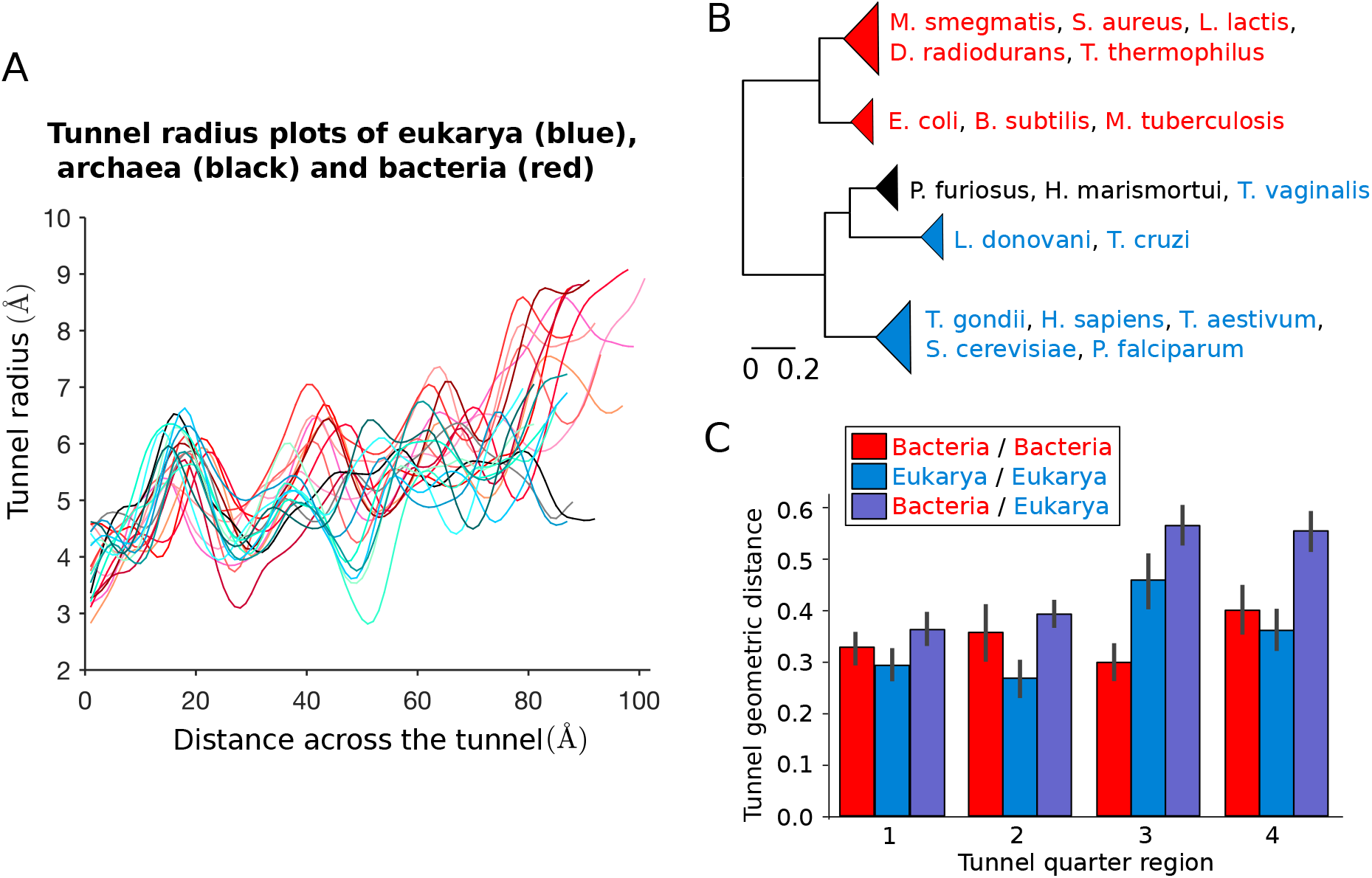
Clustering of species obtained from pairwise comparison of the tunnel geometry. A:For all our structures, we plot the tunnel radius as a function of the distance across the tunnel. These plots are used to compare the tunnel geometries (more details in Figure S1). **B:** Clustering obtained after applying our tunnel geometric distance metric to bacteria, eukarya and archaea of our dataset (see Methods). The first main branch encompasses the bacterial ribosomes, highlighted in red, while the second contains eukarya (blue) and archaea (black). For the full clustering and phylogenetic trees obtained from 16S/18S rRNA sequences, see Figure S2. **C:** We divide for each couple of species their common domain after alignment in 4 quarters (see Methods), and use the same metric to compute the distance in each of the subregions. The bar plots represent for each quarter the average and std of the geometric distance for subset of pairs made of 2 prokaryotes (red), 2 eukaryotes (blue), and 1 prokaryote and 1 eukaryote (violet).

As in the aforementioned global feature analysis, we obtained a clear separation of domains, with bacteria clustering separately from archaea and eukaryotes. Among eukaryotes, *T. vaginalis* is rather special in that it clustered with archaea ( *P. furiosus* and *H. marismortui*), which then grouped together with a cluster of trypanosomes ( *L. donovani* and *T. cruzi*); separated from these, the remaining eukaryotes formed a larger cluster. This separation of *T. vaginalis* is supported by the properties of its rRNAs, which are comparable in size to prokaryotic counterparts, and with nearly all the eukaryote-specific rRNA expansion segments missing [24]. More generally, the separations of *T. vaginalis* and trypanosomes from the other eukaryotes is also consistent with the evolutionary relationships obtained from 16S-like rRNA sequences (from the ribosome small subunit) [24]. We confirmed this result by carrying out a phylogenetic analysis for the species in our dataset, based on their 16S/18S rRNA sequences (see Figure S2A). At a finer resolution, our hierarchical clustering from tunnel geometry comparison (see Figure S2B) starts to deviate from the phylogenetic tree from sequence analysis, suggesting that smaller differences of the tunnel geometry are more difficult to relate evolutionarily. By including ribosomes from mitochondria and chloroplast (Figure S2C), we also found that they clustered with prokaryotes, which is consistent with their bacterial origin [25]. To assess the robustness of these results, we used replicate structures from same species and studied the sensitivity of the clustering to the parameters of the geometric distance (see Supporting Information, Figure S3 and Table S1). Overall, this did not lead to significant changes, demonstrating the robustness of our results to replicate structures and the choice of the comparison metric.

### The contribution of the different tunnel sub-regions to the intra-and inter-domain distance

To understand why we obtained a clear separation between the prokaryotic and eukaryotic tunnels, we more closely studied the variation of tunnel geometric distances across the two domains. Distinguishing intra-and inter-domain pairs (respectively accounting for pairs of species from the same domain, and pairs with one from each), we found that while intra-domain distances were on average similar for bacteria and eukaryotes (Figure S4), the inter-domain distances were noticeably larger (with an increase of 40% for the average). As the distance that we introduced integrates the geometric variation along the tunnel, we sought to examine which part of the tunnel contributes most significantly to the intra-and inter-domain distances. To this end, we divided up the tunnel for each pair of species into 4 quarters (the first and fourth respectively corresponding to the start and the exit parts), and applied the same metric as before to compute the geometric distance in each of these subregions (Figure 3C). While the inter-domain distance was on average always larger than the intra-domain one in every subregion, we found that across these regions, the inter-domain distance was substantially larger for the last two quarters (each carrying on average 30% of the total variation) compared to the first two ( *∼* 20% each). A similar trend was observed among eukaryotes (with 33% and 26% of the total variation respectively carried by the third and last quarters). In stark contrast, in bacteria the third quarter was the one carrying the least variation (21%, compared to 24%, 26% and 29% for the first, second and fourth subregions). Overall, our local comparative analysis of the tunnel geometry confirmed that the tunnel is more conserved in the upper part of the tunnel.

### Existence of a second constriction site in eukaryotes and the role of ribosomal protein uL4

To understand the local geometric variation of the exit tunnel observed across different species, we sought to determine how the ribosomal structure can explain the aforementioned clustering pattern of bacterial and eukaryotic tunnels. In the region associated with most inter-domain variation (see Figure 3C), we first looked at the constriction site, where proteins uL4 and uL22 meet. Interestingly, we found that the structure of uL4 at the tunnel generally differs in eukarya and bacteria, due to an extension of the uL4 loop in eukarya that yields a second constriction site. Such an extension is also present but less prominent in archaea (see Figure S5). We illustrate this structural difference in Figure 4A for *E. coli* and *H. sapiens*. The presence of a second constriction site in *H. sapiens* exit tunnel is clearly illustrated in Figure 4B, which shows that the second trough of the radius plot is even lower than the first trough for *H. sapiens*, while the opposite is true for *E. coli*.

**Figure 4.**
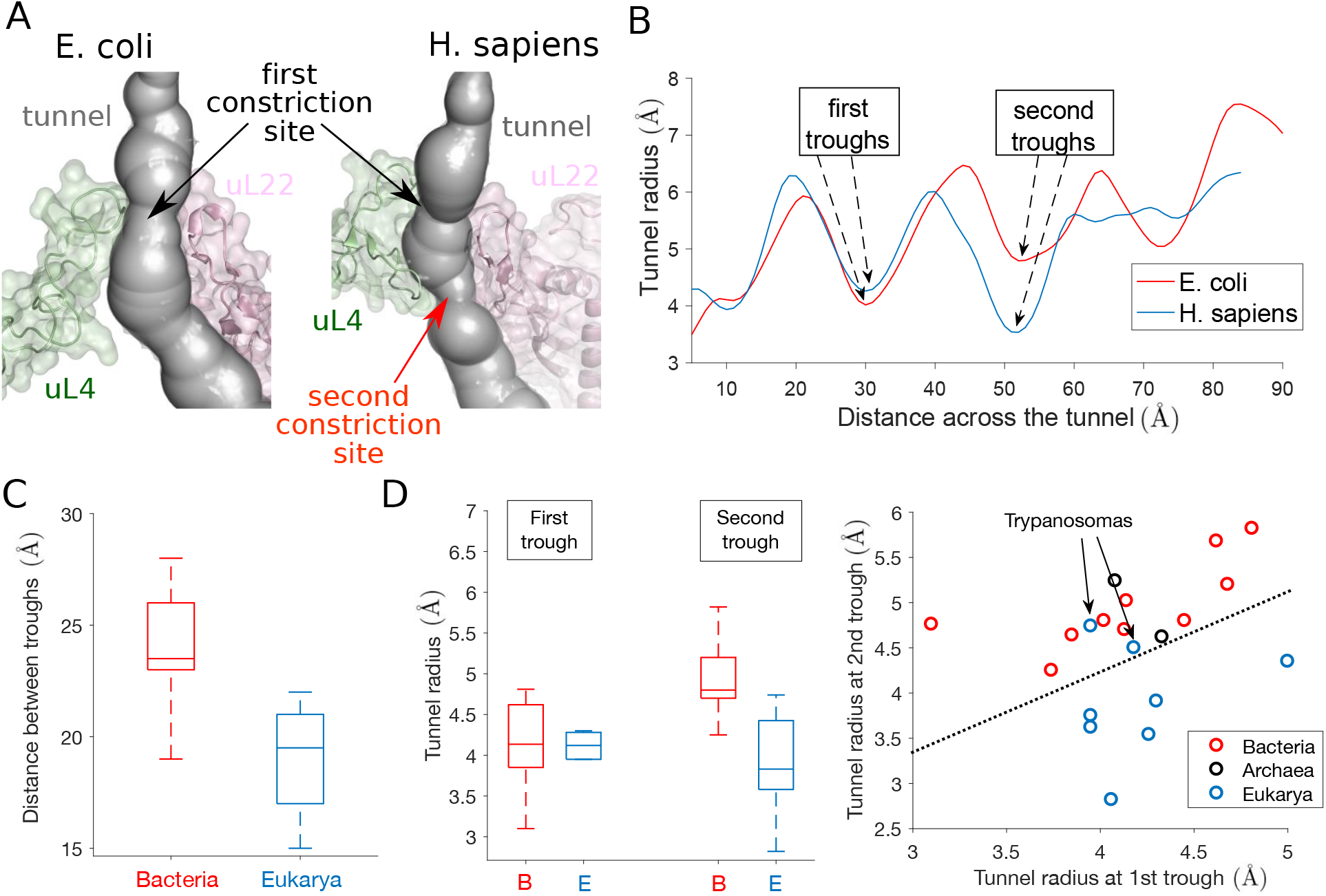
The presence of a second constriction site in eukarya explains the geometric difference observed between bacterial and eukaryotic tunnels. **A:** We show the constriction site region in *E. coli* (left), obtained from Fischer *et al.* [84] and *H. Sapiens* (right), obtained from Natchiar *et al.* [93]. The structure is surrounded by ribosomal proteins uL4 (green) and uL22 (pink). An extended arm in *H. sapiens* uL4 produces a second constriction site (see also Figure S5). **B:** The plots of the tunnel radius as a function of the tunnel distance shows a first trough associated with the constriction site (around position 30), common to *E. coli* and *H. sapiens*. A second trough appears around position 50, lower than the first trough in *H. sapiens*. **C:** We compare the distance between the troughs for bacteria (left box plot, in red) and eukarya (right box plot, in blue) in our dataset. The interquartile range is indicated by the box, the median by a line inside, and upper and lower adjacent values by whiskers. **D:** In left, we provide the same comparison as in **C** for the tunnel radii associated with the first and second troughs. In right, we compare the radius of the first and second troughs for each species of our dataset. The second trough radius is larger than the first one for all archaea (black dots) and bacteria (red dots). In contrast this is only the case in eukarya for trypanosome species *L. donovani* and *T. cruzi*.

To confirm that such a difference persists between other bacteria and eukarya, we analyzed in Figure 4C,D the relative positions and the tunnel radii associated with the first and second troughs of the radius plot for each species in our entire dataset. We found the distance between the troughs to be on average larger in bacteria (23.6 *±* 2.7Å) than in eukarya (19.1 *±* 2.3Å), suggesting structural changes in the constriction site region. While at the first trough (which corresponds to the universally shared constriction site) bacteria and eukarya have similar radii (4.2 *±* 0.6Å for bacteria and 4.2 *±* 0.4Å for eukarya), the radius at the second trough is significantly larger in bacteria (5.0 *±* 0.5Å) than in eukarya (3.9 *±* 0.6Å). Furthermore, direct comparison between the first and second troughs for each species shows that the second trough radius is larger than the first one for all archaea and bacteria. In contrast, we observed the opposite in eukarya, except for *L. donovani* and *T. cruzi* (also explaining the clustering of these two with archaea in Figure 3B). Therefore, the second constriction site in eukarya is in general narrower than the first one.

To explain the discrepancy observed for *L. donovani* and *T. cruzi*, we looked at the ribosomal structure in the constriction site region. Aside from ribosomal proteins uL4 and uL22, the tunnel is surrounded there by ribosomal RNA (rRNA). In trypanosomes like *L. donovani* and *T. cruzi*, the large subunit (LSU) rRNA breaks from the standard 28S rRNA into six smaller chains [26, 27]. The two main chains precisely meet in the constriction site region (see Figure S6), suggesting that less constraint is locally applied to the ribosomal proteins and tunnel structure, potentially increasing the size of the second constriction site. To summarize, we concluded that an important part of the geometrical differences and clustering observed between the ribosome exit tunnels come from the structure at the constriction site. More precisely, there exists a second constriction site specific to eukarya, which narrows the tunnel after the first, universally shared constriction site.

### Replacement of uL23 by eL39 ribosomal protein in eukarya affects the tunnel geometry

In addition to the structure of the constriction site region, we further examined the structure of the lower part of the tunnel, where we also detected some important geometric variations. In bacteria, the tunnel in this region is mainly surrounded by rRNA and the protein uL23. In eukarya and archaea, uL23 is also present, but the segment covering the tunnel region is replaced by the protein eL39 [19, 23, 28]. Upon close examination and comparison of these structures, we found that eL39 does not only cover the region originally occupied by uL23 in bacteria, but further extends to the tunnel exit, as illustrated in Figure 5A. More precisely (Figure 5B), the coverage distance of uL23 in bacterial ribosomes is 19.0 *±* 2.8Å, compared to 31.6 *±* 2.3Å for eL39 in eukarya. Such a difference affects the tunnel geometry (Figure 5B), as the tunnel radius in the corresponding regions is significantly larger for bacteria (6.0 *±* 0.4Å) than for eukarya (4.6 *±* 0.4Å). As a result, the exit region of the tunnel is wider in bacteria than in eukarya, with an average radius difference of approximately 1Å in the last 30Å of the tunnel (see Figure S7), hence contributing to the clustering of the radius plots that we previously obtained.

**Figure 5.**
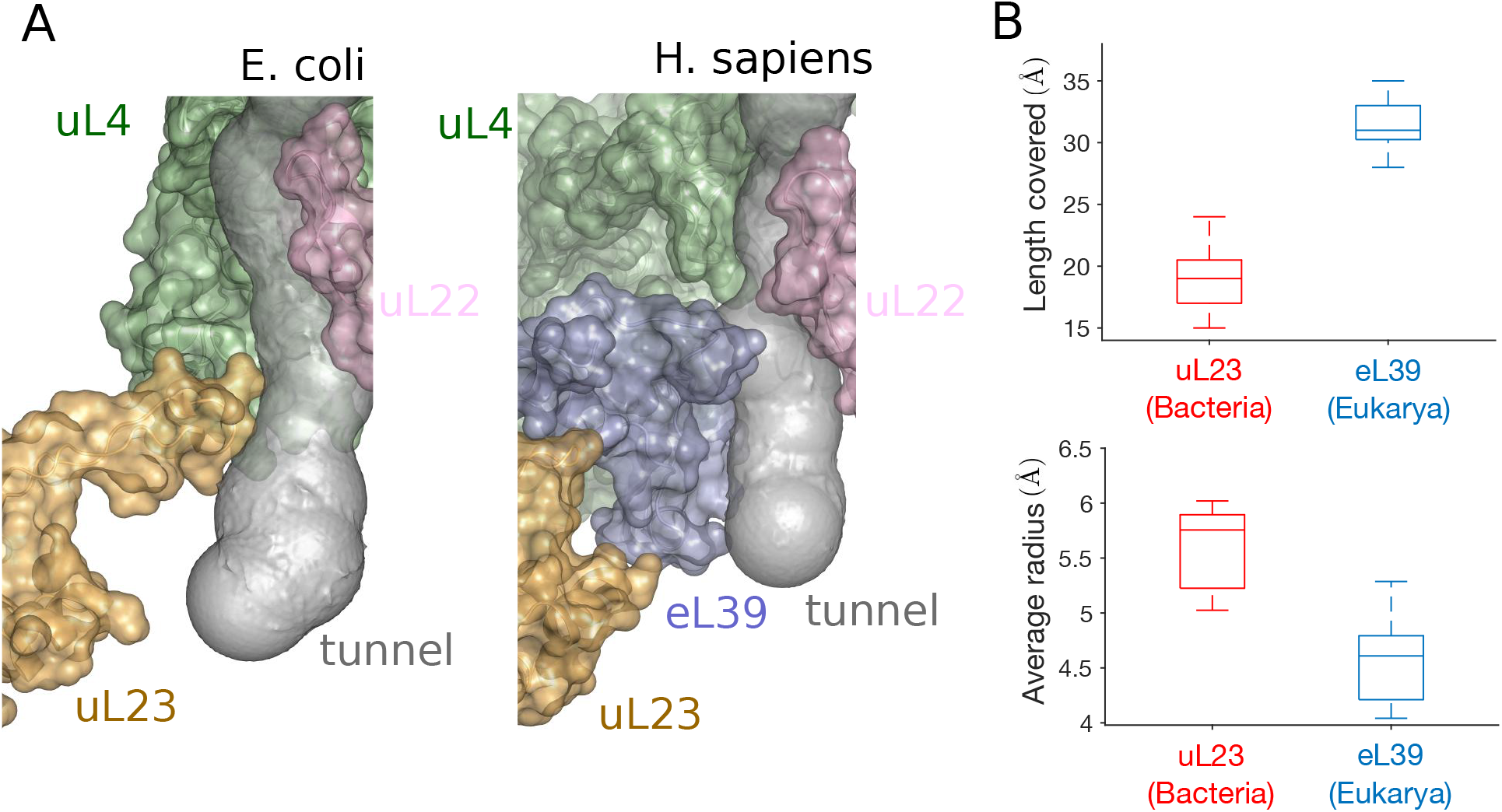
The replacement of uL23 by eL39 in eukarya impacts the tunnel geometry. **A:** The structures of the lower part of the tunnel in *E. coli* (left) and *H. sapiens* (right) show the replacement of ribosomal protein uL23 (in yellow) by el39 (in violet) in *H. sapiens*, which also covers a larger portion of the tunnel. **B:** Upper plot shows a comparison of the distance covered by uL23 and eL39 in bacteria (red box plot) and eukarya (left box plot). Lower plot shows the same comparison for the average radius.

### Association between rRNA sequence conservation and the exit tunnel

After finding evidence that the geometric variation of the tunnel across different species can be explained by rRNA and protein structural variations that have emerged through evolution, we sought to study how the tunnel and its geometry directly relate to the evolution of the ribosome at the sequence level. First, we investigated the conservation of rRNAs, focusing on the main chain (23S for bacteria, and 28S for archaea and eukarya) that constitutes the ribosome LSU. Comparative analysis of rRNA sequences has been extensively used in the past to elucidate phylogenetic relationships [29, 30], and has recently [22] led to the identification of stretches of evolutionarily conserved sequences, the so-called “conserved nucleotide elements” (CNE). In particular, it has been suggested that a large part of the CNEs may have a function in nascent polypeptide transit through the tunnel [22]. To verify this, we compared the presence of CNEs with their distance to the tunnel (see Figure 6A and Figure S8). For each species and for a given distance *d*, we computed the frequency of CNE among all rRNA nucleotides located within distance *d* from the tunnel. Plotting this frequency as a function of *d* for all the species, we observed in Figure 6B a global decrease in the frequency of CNEs as *d* increases, which means that nucleotides located farther from the tunnel are less conserved. For much divergent rRNAs such as the ones from organelles or *T. vaginalis* [24], the low number of CNEs yields no association with the distance to the tunnel. For bacteria, archaea, and eukaryotic trypanosomes, we found that the frequency of CNEs within 25Å from the tunnel is 21 *±* 5%, which is more than the double the frequency of CNEs located between 25 and 50Å (10 *±* 2%) and also that located farther than 50Å from the tunnel (8 *±* 2%). In the remaining eukaryotes, we found a much larger frequency of CNEs within 25Å from the tunnel (45 *±* 12%), with a sharp decrease for the region between 25 and 50Å (18 *±* 5%) and the region beyond 50Å (10 *±* 3%). Distinguishing universal and domain-specific CNEs [22], we found a similar trend for both, with a larger contribution from domain-specific CNEs (Figure S9). Overall, these results confirm the association between the tunnel and the conservation of surrounding rRNA nucleotides.

**Figure 6.**
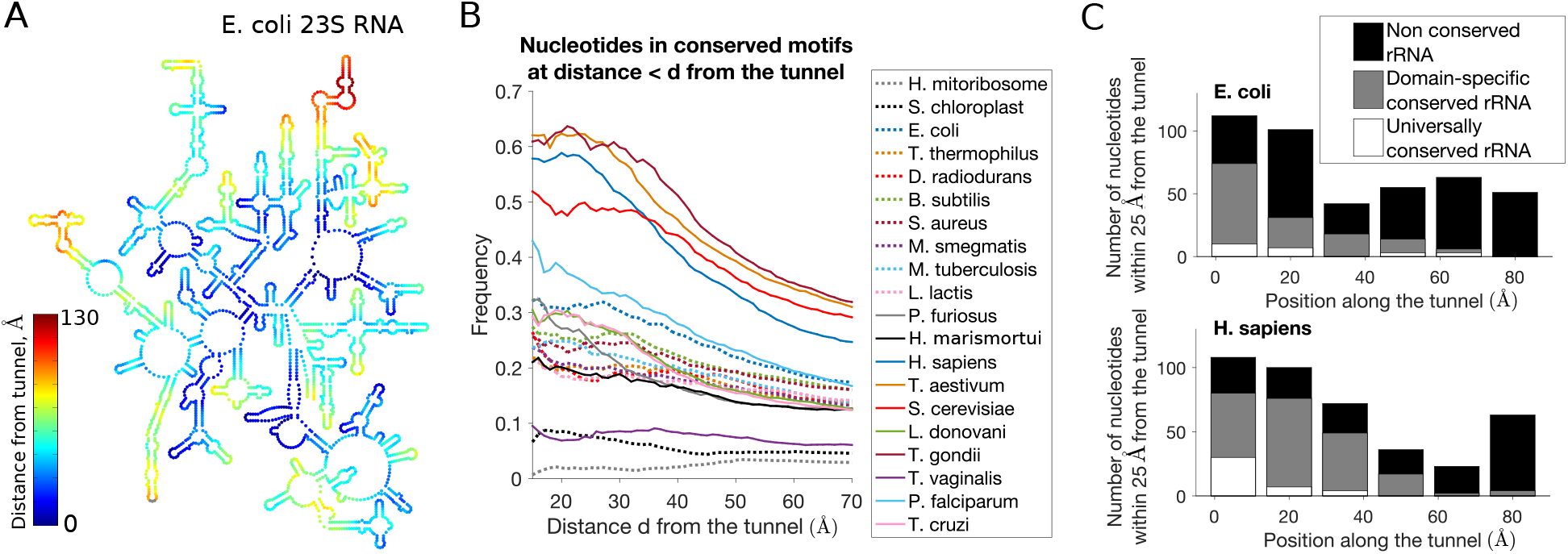
Association between geometric and sequence conservations of ribosomal rRNA. **A:** A map of the secondary structure of the 23S rRNA in *E. coli*, colored by the distance from tunnel (see also Figure S8). **B:** For a given species and distance *d*, we look at all the rRNA nucleotides located within distance *d* from the tunnel, and we compute the frequency of conserved elements [22]. We plot this frequency as a function of *d* for all the species of our dataset (see also Figure S9). **C:** We study the local conservation of rRNA nucleotides along the tunnel: Upon dividing the tunnel into regions of 15Å along the centerline, we consider for each region all the rRNA nucleotides that are the closest and located within 25Å, and we compute the associated number of conserved, domain-specific and universally conserved elements. We show here the resulting plots for *E. coli* (up) and *H. sapiens* (down) (for other species, see Figure S10).

To determine whether such conservation is homogeneous or specific to some local parts of the tunnel, we computed for each species the local frequency of CNEs and compared the level of conservation across the tunnel, as shown in Figure 6C and Figure S10. Overall, we found that the sequence conservation along the tunnel is heterogeneous and strongly enhanced towards the upper part of the tunnel, as more than 80% of the CNEs were associated with the first 40Å of the tunnel (while all the nucleotides positioned along the first 40Å of the tunnel represent only 48% of all nucleotides). In this region, we also observed a stronger conservation closer to the PTC, as 43% of the CNEs were located less than 10Å from the tunnel start position. These results are in agreement with the comparative geometric study of the tunnels, which also showed more conservation in the upper part region.

### Conservation of ribosomal protein sequence and positive charges at the exit tunnel

Lastly, we studied the relation between the geometric and sequence conservations at the tunnel for ribosomal proteins. To do so, we aligned across species the main proteins located close to the tunnel, namely uL4, uL22, and uL23 for bacteria and eL39 for eukarya (see Methods, Figure 7A and Figure S11). For uL4, because of multiple insertions that prevent a good alignment of all sequences (like the one leading to the aforementioned second constriction site), we separately aligned uL4 for bacterial and eukaryotic sequences. Upon computing a conservation score [31] for each alignment, we found peaks of conservation in regions located in proximity with the ribosomal tunnel, suggesting that association between the sequence conservation and the exit tunnel also occurs for proteins (Figure 7B and Figure S12A). Upon closer examination of the consensus motif sequences at the tunnel, we found a large fraction ( *∼* 30%) of positively charged amino acids (arginine R and lysine K). Interestingly, these positively charged amino acids sometimes co-occur at the same position of an alignment, e.g., position 159 of the uL22 alignment (with 8 arginines and 8 lysines, see Figure 7A), position 69 of bacterial uL4 alignment, and position 34 of eL39 alignment (Figure S11). This suggests that not only the sequence but also the charge properties are important to maintain the integrity of the tunnel.

**Figure 7.**
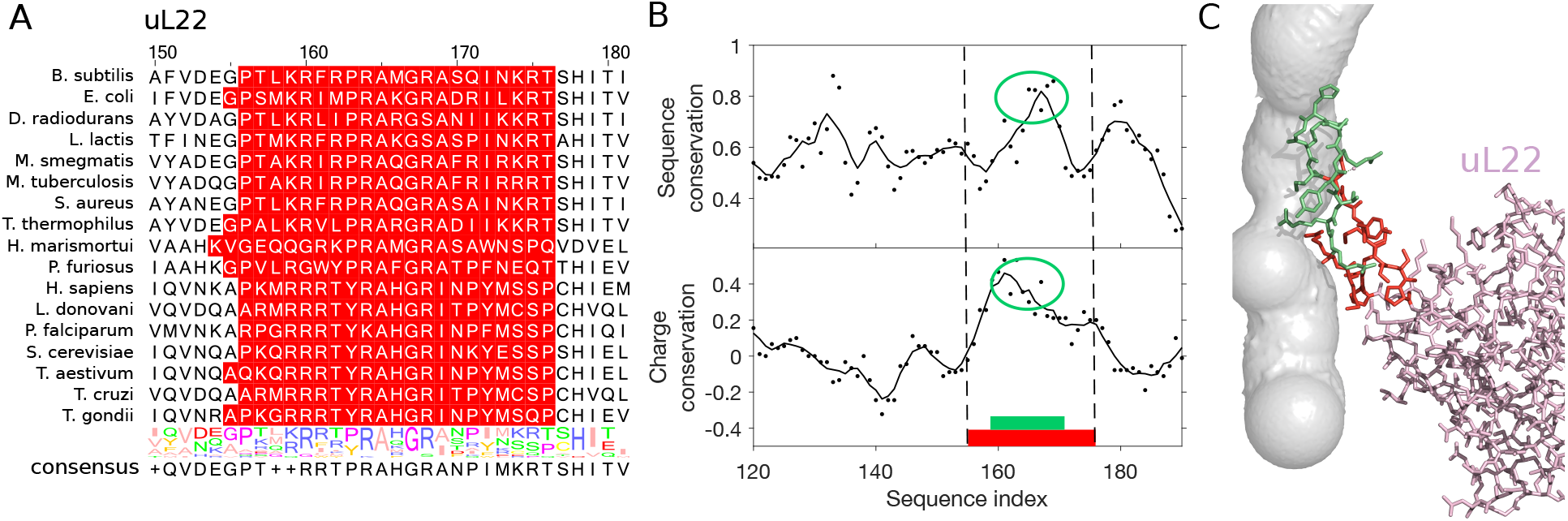
Conservation of sequence and positive charge of ribosomal protein uL22 at the tunnel. **A:** We show the multiple sequence alignment of ribosomal protein uL22 close to the tunnel. Residues highlighted in red are the ones located within 10Å from the tunnel. **B:** We plot the sequence and charge conservation scores (see Methods) along the sequence alignment. Continuous lines represent the signal averaged over a window of 5 sites. Region in red is the same as in **A**. In green, we highlight a subregion of residues close to the tunnel, with a peak in charge or sequence conservation. **C:** The associated structure of uL22 in *H. sapiens*, where residues in green and red correspond to the ones highlighted in **B**. In particular, the green region of high charge and sequence conservation is also in direct contact with the constriction sites. For the other ribosomal proteins associated with the tunnel, see Figure S11 and Figure S12.

Therefore, we introduced a measure of charge conservation (see Methods) that specifically accounts for the local presence of positively charged amino acids in an alignment. Recent studies have shown enrichment of positively charged amino acids in ribosomal proteins [32]. Upon computing the charge score for our sequence alignments, we indeed found some significant correlation between the sequence conservation and charge conservation scores for bacterial proteins with Pearson’s correlation *r*^2^ = 0.70, 0.51 and 0.47 (p-values < 10^−4^) for bacterial alignments of uL4, uL23 and uL22, respectively (see Supplementary Data). We also found that overall, regions with the highest presence of positive charges are located in the tunnel region (see Figure 7B and Figure S12A). In uL22, the residues in direct contact with the tunnel (Figure 7B and C) are highly conserved, with the highest charge conservation score. Larger charge and sequence conservations were also found in uL4, with more conservation at the first constriction site compared to the second one in eukarya (Figure S12A and B). In the lower part of the tunnel, we detected a strong peak of both charge and sequence conservation in bacterial uL23. Interestingly, while we did not observe in eukarya a conservation signal as strong in the whole region of the tunnel covered by eL39, the region where eL39 overlaps with uL23 in bacteria also exhibits large charge and sequence conservation (Figure S12A and B), suggesting persistence of the local charge properties in this region although the ribosome structure differs between bacteria and eukarya. Overall, we concluded that ribosomal proteins are more locally conserved at the proximity of the tunnel, with a bias towards positively charged amino acids, suggesting that these amino acids and the electrostatic environment are important for the tunnel function (see Discussion).

## DISCUSSION

In this study, we carried out a comparative analysis of the ribosome exit tunnel geometry that is most comprehensive to date. Recent advances in cryo-EM [33, 34] allowed us to observe the ribosome at a resolution fine enough to characterize the local structure of the exit tunnel, thereby making it feasible to detect significant differences across species. Despite the eukaryotic 80S ribosome being approximately 50% greater in mass and rRNA lengths compared to the prokaryotic 70S ribosome [21, 35], we found the bacterial exit tunnel to be larger, in both length and average radius (Figure 2). Such differences mostly come from the lower part of the tunnel.

We classified the tunnels using their radius variation plots and interestingly found that the resulting hierarchical clustering closely reflects the species phylogeny (Figure 3). We then found evidence of two notable evolutionary changes that affect the tunnel structure and explain our clustering result. The first is the insertion of an extended loop sequence in eukaryotic ribosomal protein uL4 (Figure 4), which leads to a second constriction site that is narrower than the first one (except in trypanosomes). The second is the replacement of the uL23 loop in the lower part of the tunnel in bacteria by the small protein eL39 in archaea and eukarya (Figure 5). While the existence of the first constriction site and the overlapping positions of eukaryotic eL39 and prokaryotic uL23 proteins have been previously well established [23, 28, 36], our comparative analysis shows that the above evolutionary changes have led to important differences among species that significantly affect the tunnel geometry and hence the path of the nascent chain through the tunnel.

### Implications on the evolution of the ribosome

Since both ribosomal RNA and protein sequences located near the exit tunnel are more conserved (Figure 6, Figure 7, and Figure S12), there is a strong relationship between the evolution of ribosome components and the variation in the geometry of the exit tunnel [22]. The high geometric conservation of the upper part of the tunnel coincides with the high sequence conservation of the associated rRNAs and ribosomal proteins, suggesting that it is an important part of the ribosome core. The observed contrast in conservation between the upper and lower parts of the tunnel is in agreement with previously proposed models of the evolution of the rRNA structure [21], which distinguishes an early phase of creation of a short tunnel, subsequently completed by a phase of tunnel extension and expansion of the LSU. These two phases are preceded by the maturation of the PTC, commonly considered as the oldest ribozyme [21, 37, 38] and which also appeared in our analysis as the most universally conserved region [22, 39]. While there is a clear association between the geometry of the exit tunnel and the evolution of its components, some other biophysical properties of the tunnel may also be influencing the evolution of the ribosome. Upon closer examination of the conserved ribosomal protein sequences at the tunnel, we indeed found a significant contribution of positively charged amino acids (see Figure 7 and Figure S12), which are generally found to be highly enriched in ribosomal proteins [32, 40]. While it has been proposed that this general enrichment may be needed to create electrostatic interaction during ribosome assembly [40], our analysis suggests that the conservation of positively charged amino acids near the tunnel could be tied to maintaining an appropriate electrostatic environment inside the tunnel (see below).

### Impact on the nascent polypeptide chain transit

The geometric and structural variation of the exit tunnel may have several functional and biophysical implications, as the nascent chain transit can be affected. On the one hand, a too narrow tunnel could obstruct the elongation of the nascent chain and generate tunnel-peptide interactions that might alter the elongation rate during protein synthesis [4, 41, 42]. On the other hand, a too large tunnel radius could lead to an increase of conformational sampling [43, 44, 45] and hence misfolding events. Our results indicate that before reaching the constriction site in the upper tunnel, the nascent chain propagates in an environment that seems relatively well conserved, with an increase of the tunnel radius followed by a decrease near the constriction site. In the context of a diffusing particle inside the tunnel, we previously found [9] that this radial increase creates a strong entropic barrier, which can be compensated by the electrostatic potential if the particle is positively charged. More generally, it was also experimentally shown that electrostatics in the ribosomal tunnel modulate chain elongation rates [9, 17] and can even induce ribosome stalling [46]. At the proteome-wide level, averaging the charged amino acid frequency over all translated genes in *S. cerevisiae* revealed an elevated (respectively, decreased) amount of positively (respectively, negatively) charged amino acids in the first ∼ 20 codons [9]. These patterns also hold in other species (Figure S13). Together with the conservation of ribosomal protein charge at the tunnel (which is also more significant at the first than at the second constriction site, see Figure 7 and Figure S12), they suggest co-evolution of the tunnel and the charge distributions of the proteome, to maintain a favorable electrostatic environment for the nascent polypeptide during its initial pass through the exit tunnel.

Once the N-terminus exits from the tunnel, the elongation of the nascent chain may be affected by additional factors, such as the co-translational processes that generate pulling forces on the nascent polypeptide chain [47]. For example, the translocon, involved in the translocation of membrane proteins, can relieve elongation arrest due to the SecM sequence, thereby enabling translation restart [48]. In eukaryotes, a similar mechanism has been suggested for the chaperone Hsp70 [49]. More generally, proteins that start to fold co-translationally while still in contact with the ribosome, exert some pulling force sufficient to weaken or even abolish stalling [50, 51]. As shown by theoretical studies of polymer dynamics in response to external pulling [52, 53], the efficiency of the pulling depends on how tension propagates from the ends into the bulk, and, in our case, how the free energy released by the folding reaction can get stored as an increased tension in the nascent chain [54]. This phenomenon is evidenced by the increase of the pulling force with the nascent chain hydrophobicity (found with the formation of *α*-helix structure and chain stiffness [47]) and the positive impact of the hydrophobicity on the elongation rate (obtained by analyzing ribosome profiling data [9]). As the confined geometry of the tunnel can actually play a role in stabilizing *α*-helices [55], we therefore hypothesize that the presence of a second narrower constriction site in eukaryotes can improve the tension propagation across the nascent chain, and thus facilitate the elongation process.

### Functional implications of the tunnel variation

We detected more variation in the lower part of the tunnel, as structural modifications of the ribosomal proteins lead to a decrease of the eukaryotic tunnel size. Various experiments and simulations have shown that the tunnel can be large enough to accommodate a substantial degree of protein structure (e.g., *α*-helix, tertiary hairpins and even protein folding), especially in the so-called “folding vestibule” located near the exit port [36, 54, 56, 57, 58, 59]. As the shape of the exit tunnel in this region can affect co-translational protein folding [60], our results suggest that eukaryotic tunnels are less favorable for such folding to occur inside the ribosome. Such a difference could actually reflect the differences in complexity and division of labor between the eukaryotic and prokaryotic chaperone networks [13, 56]. In prokaryotes, both co-translational folding and denaturation of proteins during stress are ensured by an overlapping set of chaperones that primarily relies on the bacterial trigger factor [61]. In contrast, distinct specialized networks of downstream chaperones evolved in eukaryotes to separately carry out these processes [62]. In particular, the existence of a more efficient chaperone network devoted to folding may be the result of eukaryotes having a higher proportion of larger multidomain proteins, as well as more complex protein folds [63]. While the prokaryotic tunnel can assist the chaperone network by pre-folding the nascent chain inside the tunnel [64], the Prefoldin family of proteins [65] plays such a role instead in eukarya and archaea, which could explain the observed reduced size of the tunnel for these domains.

Another example of the direct implication of the bacterial ribosome tunnel on co-translational processes is the translocation of membrane proteins, which is mediated in both eukarya and bacteria by the Signal Recognition Particle (SRP), a ribonucleoprotein that recognizes the signal peptide emerging from the ribosome [1, 8, 66, 67]. Such a process is constrained by the short time window to recognize and target the signal peptide [66, 68]. In bacteria, the SRP can be recruited prior to the emergence of the peptide from the tunnel [69], via uL23 that both recognizes the nascent sequence inside the tunnel and binds to the SRP at the outer surface of the ribosome. In eukaryotes, however, forward signaling cannot similarly occur, as the replacement of uL23 by eL39 disrupts the original coupling between the signal peptide inside the tunnel and the SRP recruitment [8]. To compensate for this lacking function, it has been shown that eukaryote-specific signaling mechanisms can promote the progressive binding of the SRP [68], while the usage of non-optimal codons, associated with translation slowdown, also increases the binding opportunity time window for the SRP [66].

Finally, while the reduction of the tunnel size in eukarya seems to reflect the externalization of major co-translational functions, the increase of tunnel confinement may also provide some other advantages. Besides the aforementioned facilitation of nascent chain elongation, the reduction of the tunnel radius at the exit port by eL39 and the addition of a second constriction site also contribute to restricting the access to the tunnel and the PTC from external threats. For example, as the insertion of amino acids in the loop of uL4 in *E. coli* confers resistance to larger-size macrolides [70, 71], it has been suggested that the narrower size of the constriction site in eukarya can block the access of these antibiotics to the targeted PTC; we actually found a second eukaryote-specific constriction site, located below the universal constriction site, to be responsible for this narrower access. Similarly, mutants lacking eL39 are more vulnerable against ribosome targeting antibiotics [72], in addition to an increased translation error rate and cold sensitive phenotype. Molecular dynamic simulations have predicted interaction between the 28S rRNA tetraloop and eL39, potentially leading to even more obstruction of the tunnel, suggesting that eL39 acts as an energy barrier to ensure protein quality control and protect the ribosome from deleterious external agents [19].

### Future directions

Based on the present study and the aforementioned involvement of the tunnel structure in various co-translational processes, it would be interesting to thoroughly investigate the inter-domain differences in the chaperone machinery engaging the ribosome exit tunnel, especially near the exit port. For example, the class of chaperones involved in guiding *de novo* protein folding includes multiple domain-specific actors, such as the bacterial trigger factor, the archaeal and eukaryotic nascent polypeptide-associated complex and specialized eukaryotic heat shock proteins [61], the diversity of which may be the result of different strategies developed in conjunction with the tunnel structure. Elucidating the evolutionary basis of these processes might also require the development of more refined tools to extract and analyze the tunnel geometry and its properties. It is quite remarkable that only considering the radius variation and a simple metric allowed us to recapitulate the general phylogenetic relationship. Integrating more information from the geometry and structure of the tunnel should allow to study and interpret smaller variations of the tunnel, which should also become more reliable as the resolution of the structures improves. In particular, it would be interesting to build a stochastic model describing the evolution of the tunnel geometry, to decipher the larger variation detected in bacteria and to determine if it is mainly due to the difference in evolutionary time scale or whether natural selection played a role.

As the tunnel wall responds to the nascent peptide contained within it [73], it would be important to analyze and compare the geometry of the tunnel in the presence of a nascent peptide, and specifically arrest sequences known to interact with the tunnel. While the responsiveness of the tunnel to the chain is in general likely to be at the level of small movements [74], we interestingly found that the tunnel structure of *E. coli* containing the ErmCL chain seemed to deviate more substantially from the original one in the region where the sequence spans (Figure S3). The methods presented here could be extended and applied to more structures in the future, and as a result provide useful insights into many of the essential biological processes in which the ribosome takes part.

## METHODS

### Ribosome structures

Cryo-EM reconstructions and X-ray crystallography structures of ribosomes were downloaded from the protein data bank https://www.rcsb.org/. References and details are given in Table 1. The fitting of the model’s residues to the original map was evaluated for each structure using coot [75] density-fit-score function (2Fo-Fc maps were used for X-Ray data).

### Extraction of the ribosome tunnel geometry

To extract the tunnel coordinates, we used a tunnel search algorithm developed by Sehnal *et al.* [76]. The tunnel search was initiated at the PTC. To locate the PTC, we aligned the sequences of 23S and 28S rRNAs and selected the nucleotide aligned with U4452 in human. The tunnel search algorithm was applied after editing the structure to contain atoms located less than 80Å from the constriction site. For the human ribosome, the constriction site was obtained by computing the center of mass of amino acids G70, R71, in uL4 and H133 in uL22. The same procedure was applied for the other species after alignment of proteins uL4 and uL22. As the algorithm gives several possible tunnels, we manually picked the ribosome exit tunnel by looking at the shape, length and relative position to uL4 and uL22. Coordinates were extracted using Pymol and Python custom scripts. The origin of the tunnel was set at a distance of 5Å from the nucleotide used to initiate the tunnel search algorithm, to remove sensitive regions generated by the tunnel search algorithm close to the initial point. Downstream analysis for computing global and local geometric features of the tunnel were done using Matlab and Python (More details in Supporting Information).

### Distance metric for pairwise comparison and clustering of radius plots

For two tunnels *T*_1_ and *T*_2_ parametrized by 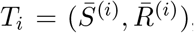, where *i* = 1, 2, 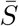 is an arc length parametrization of the tunnel centerline 3D coordinates and 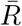 is the associated radius (in ångström), we introduced the following distance metric *D* (*T*_1_*, T*_2_) given by

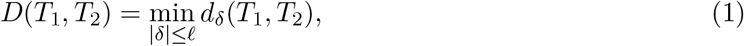

where 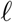 is the maximum shift length and

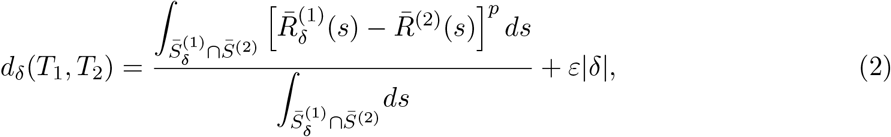

where 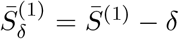, and 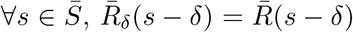. In other words, we looked for the best alignment of the two tunnels such that it minimizes the average *L^p^* difference between the two aligned radius plots, with a negative penalty for the size of the alignment shift (which is also limited to be less than 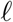). We then take this measurement as the distance between the two tunnels. In practice, we took 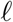 = 20Å, *ε* = 0.01 and *p* = 2 (for more details on how sensitive our results are with these parameters, see Supporting Information), and evaluated the integrals by computing their Riemann sum. After computing the pairwise distance matrix associated with our dataset, we constructed the associated phylogenetic tree using the Unweighted Pair Group Method Average (UPGMA) algorithm.

### Sequence alignment and conservation of ribosomal proteins

Ribosomal proteins are named according to the system set by Ban *et al.* [77]. Sequences were aligned using MAFFT [78] with default parameters and visualized in Jalview [79]. Conservation scores were computed using Jensen-Shannon divergence and software provided by Capra and Singh [31], using the default parameters.

### Conserved motifs in ribosomal RNA

For each of the three domains (bacteria, archaea and eukarya), conserved motif sequences were obtained from [22], with their positions along the rRNA sequence computed using exact string matching. For each nucleotide, the distance from the tunnel (or PTC) was computed using MATLAB.

### Charge conservation score

To quantify charge conservation among the ribosomal proteins in our study, we introduce the following metric: For a given multiple sequence alignment (MSA) *M* containing *N* sequences, let *MC* denote the *C^th^* column of the MSA and 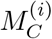 denote the symbol in column *C* of sequence *i*, such that 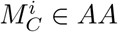, where *AA* is the set of 21 amino acids plus the gap symbol. For each element *x* of *AA*, we assign a charge *c*(*x*) = +1 for lysine and arginine, −1 for aspartic acid and glutamic acid and 0 for all other amino acids plus the gap symbol. For each column *C* of the MSA, we define the conservation charge score *S*(*C*), as

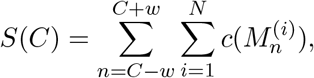

where the window size *w* is set in our analysis to be 2. Analysis of conservation scores was done using Matlab custom scripts.

### Visualization tools

Structures were visualized using Pymol. Maps of ribosomal RNA secondary structures, distance of the rRNA nucleotides to the tunnel (or PTC) and conserved motifs were visualized in RiboVision [80].

## Acknowledgments

This research is supported in part by NIH grants R01-GM065050 and R01-GM094402, and a Packard Fellowship for Science and Engineering. YSS is a Chan Zuckerberg Biohub investigator.

